# Machine learning-based modulation of Ca^2+^-binding affinity in EF-hand proteins and comparative structural insights into site-specific cooperative binding

**DOI:** 10.1101/2021.10.04.462978

**Authors:** Mohit Mazumder, Sanjeev Kumar, Devbrat Kumar, Alok bhattacharya, S. Gourinath

**Affiliations:** School of Life Sciences, Jawaharlal Nehru University, New Delhi-110067; Department of Biochemistry and Molecular Biology, Michigan State University, East Lansing, Michigan 48824, USA

**Keywords:** Ca^2+^-binding affinity, EF-hand proteins, protein engineering, Isothermal calorimetry, X-ray crystallography, support vector machine, dynamic residue correlation

## Abstract

Ca^2+^-binding proteins are present in almost all living organisms and different types display different levels of binding affinities for the cation. Here, we applied two new scoring schemes enabling the user to manipulate the binding affinities of such proteins. We specifically designed a unique EF-hand loop capable of binding calcium with high affinity by altering five residues of the loop based on the scoring scheme. We worked on the N-terminal domain of *Entamoeba histolytica* calcium-binding protein1 (Nt*Eh*CaBP1), and used site-directed mutagenesis to incorporate the designed loop sequence into the second EF hand motif of this protein. The binding isotherms calculated using ITC calorimetry showed a ∼500-fold greater association constant (K_a_) for the mutant. The crystal structure of the mutant was also determined, and displayed more compact Ca^2+^-coordination spheres in both of its EF loops than did the structure of the wildtype protein, consistent with the greater calcium-binding affinities of the mutant. The Nt*Eh*CaBP1 mutant was also shown to form a hexamer rather than just a trimer, and this hexamer formation was attributed to the position of the last helix of the mutant having been changed as a result of the strong calcium coordination. Further dynamic correlation analysis revealed that the mutation in the second EF loop changed the entire residue network of the monomer, resulting in a stronger coordination of Ca^2+^ even in other EF hand loop.

## Introduction

Ca^2+^ is a ubiquitous secondary messenger that controls various cellular processes like cell differentiation, muscle contraction, exocytosis, and apoptosis event [1, 2]. One of the well-studied Ca^2+^ binding proteins is Calmodulin (CaM), known to interact with many cellular proteins thereby performing various functions [3-5]. The CaM or CaM-like proteins has a conserved Ca^2+^ binding motifs referred as EF-hand motifs. Including CaM and other CaM like proteins, the number of EF-hand motifs varies across the different CaBPs [6-8]. The EF-hand containing proteins are abundantly present in the living organism and are classified more than 66 subfamilies and plays numerous roles in many crucial pathways [5].

The residues of EF-hand motifs are well conserved across the various species, and mostly by negatively charged amino acids. The coordination of amino acid residues of EF-hand motifs with Ca^2+^ occurred in a very conserved manner that forms a pentagonal bipyramidal geometry [5, 7, 9, 10]. In most of the Ca^2+^ binding proteins, including all CaMs, the residues in the Ca^2+^ binding loop at positions +X(1^st^ residue of the loop), +Y(3^rd^ residue of the loop), +Z(5^th^ residue of the loop), - Y(7^th^ residue of the loop), -Z (9^th^ residue of the loop) and -X (12^th^ residue of the loop) coordinate with Ca^2+^ directly, the 9^th^ (*-Z*) position residue does not coordinate the Ca^2+^ directly, instead a water molecule involved in the coordination and 12^th^ position residue contributes donates two electron and complete the coordination sphere [7, 9-12].

Binding of Ca^2+^ to CaM/CaM-like proteins, induces the conformational rearrangement which changes the energy landscape of these proteins, thereby interacts with many intracellular proteins [13]. The residue composition in the EF-hand binding loop is one of the major determinants, which dictate the Ca^2+^ binding affinity of all CaBPs; other factors such as the conformational cost upon Ca^2+^ binding, the EF-β-scaffold, cooperative binding, the physiological environment, also play crucial roles in the binding of the metal ion [5, 12, 14, 15]. These factors lead to a wide range of Ca^2+^ binding affinities, perhaps such diverse Ca^2+^ binding affinity exhibited by CaBPs may allow them to perform variety of functions.

Due to the importance of Ca^2+^ signaling, the Ca^2+^ binding proteins have been extensively studied; for example, various mutational studies have been reported on the Ca^2+^-binding loops of CaM [16, 17]. Many site-specific binding affinity studies, using methods such as grafting the EF-hand loop into different proteins such as CD2 proteins, have been conducted to understand the role EF-hand residues in Ca^2+^ binding [18-22].

Previously, we developed a method to identify and classify the Ca^2+^-binding EF-hand loops by levels of their Ca^2+^-binding affinities using support vector machine (SVM) based approaches operating through a classification-based scoring system [23]. The classifier uses a nonlinear SVM with a gaussian radial basis function kernel [24]. The program showed high accuracy for the prediction of Ca^2+^ binding motifs available from the literature but it only predicts the affinities qualitatively. Despite of huge implication of EF-hand motifs mediated Ca^2+^ signaling across the various species, there is no proper method has been developed yet which helps in manipulating Ca^2+^ binding affinity systematically. Although Ca^2+^ binding affinity can be manipulated by mutating critical residues of EF-hand motifs (1^st^,3^rd^, 5^th^, 7^th^ and 12^th^) of EF-hand motifs, but it often abolishes the Ca^2+^ binding affinity completely, which limits the study the function of CaBPs over a wide range of Ca^2+^ binding affinity.

To overcome this shortcoming in the field of Ca^2+^ signaling, we upgraded our previously available software in a way so that user can not only predict the Ca^2+^ binding affinity qualitatively rather one can manipulate the binding affinities *in-vitro* by mutating the EF-hand motif residues. For upgrading our current software, we extracted the margin distances for each prediction from our non-linear hyperplane which is the distance between the decision boundary and the support vectors and calculated the weighted position-specific scores (PSM) for each prediction [23]. With two additional parameters to further classify the EF-hand loops, we designed a unique EF-hand loop with predicted capability to bind Ca^2+^ with high affinity. The machine trained to identify Ca^2+^ binding sites & estimate affinity was further used to design and discriminate between two similar sites based on new scores.

To validate the binding abilities of the new designed site, we introduced the designed sequence in N-terminal domain of *Entamoeba histolytica* Ca^2+^ binding protein1, EF-hand loop 2 (Nt-EhCaBP1 EF-2 mutant) and we applied, biochemical, structural, and computational approach to validate our hypothesis. The binding affinity determined using ITC clearly shows Nt-EhCaBP1 EF-2 mutant forms of protein binds Ca^2+^ with higher affinity than Wild type-Nt-EhCaBP1 (Wt-Nt-EhCaBP1). Furthermore, we employed structural biology approach (X-ray crystallography) to understand the changes at the atomic level, that lead to changes in its Ca^2+^ binding affinity. Further, we used normal mode analysis (NMA) [25] to obtain insights leading into the coupled motions and allosteric effects directly influencing the interactions in two Ca^2+^-binding sites of each protein. The analysis indicated a direct influence of the EF-hand loops on the central helix as a result of long and short-range interactions affecting the entire residue network and hence enhancing the allosteric interactions in high-affinity sites. Based on sequence and structural investigations followed by the network analysis, we derived the basis of allosteric cooperativity shown by different EF-hand Ca^2+^-binding loops.

A set of scripts with new scoring functions and a user-friendly webserver to predict, design and engineer EF-hand binding loops were developed. The webserver and the downloadable set of scripts are available at http://sbl.jnu.ac.in/calb/index.html.

## Materials and Methods

### Scoring system for each prediction

Previously, we developed a method to identify and classify Ca^2+^-binding EF-hand loops [23]. The classifier uses support vector machine (SVM) machine learning with C and Gamma parameters for specifically a nonlinear SVM with a Gaussian radial basis function (RBF) kernel [23]. To increase the efficiency of the classification, we extracted the margin distance between the decision boundary and the support vectors for each prediction.

In general, margin is referred to as the distance of the vector (x) which is the Euclidean distance of x from the separating hyperplane [26], is given by

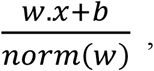

Where w = perpendicular to the hyperplane and b is, the parameter used to determine the offset of the hyperplane from the origin along the normal vector.

The distance to the origin was calculated as 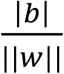. The margin (d (margin)) was calculated using the equation*d*(*margin*)=*m*_*positive*+*m*_*negative*. The output is a signed distance that is either negative/positive, depending on which side of the hyperplane the point x resides [27, 28].

### PSSM based log likelihood scores (PSM_LogL_)

The conservation score based on PSSM (position specific scoring matrix) for each sequence submitted for classification was calculated by using the equation *Gij*=*log*(*Sij*/*Pi*).Where *Gij* is the calculated PSSM score and *Bij* is the frequency of amino acid residue *i* at position *j* in matrix *S*, and was calculated by using the equation

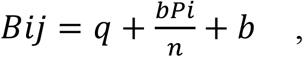

where q denotes the observed counts of amino acid residue type i at position j, *Pi* is probability of amino acid residue type i, b is a pseudocount considered here as the square root of the total number of training sequences, and n is the number of training sequences. *Gij* (PSSM score) represents the foreground model, which is the true homology and *Pi* represents the probability that a match occurs at random (background model) calculated using the BLOSUM62 substitution matrix [29]. In short, the PSSM-based scoring includes the relative frequencies obtained by counting the number of times each amino acid residue occurred at each position of the alignment, followed by normalization of the frequencies.

### Design of the unique EF-loop site

We used SVM margin score (SVM_Mar_) and the PSSM-based log likelihood score (PSM_LogL_) values to design a unique Ca^2+^-binding site, one does not present in any of the protein sequence databases [30, 31]. The design was developed in order to improve the predictions of the classifier as well as to understand the binding mechanism by deciphering the roles of Ca^2+^-binding residues. The sequence design was carried out by substituting each of the 20 encoded amino acid residues in all possible 12 positions in the Ca^2+^-binding loop; giving a unique score with every iteration. The sequence DKDGDGFIDFEE showed a high score and the proteome wide BlastP [30] showed the absence of the site in all known databases. Therefore, the 2^nd^ EF-hand loop of Nt*Eh*CaBP1, i.e., DADGNGEIDQNE, was replaced with the DKDGDGFIDFEE sequence. This substitution was carried out using site-directed mutagenesis. Note that Nt*Eh*CaBP1 denotes the N-terminal domain of the EF-hand-containing Ca^2+^-binding protein1, which has been well characterized in our laboratory [32-34]. The N-terminal domain has two Ca^2+^-binding EF hand motifs. In order to construct the desired mutant, we incorporated 5-point mutations, namely A47K, N50D, E52F, Q55F and N56E, in the 2^nd^ EF-loop of Nt*Eh*CaBP1.

We selected Nt*Eh*CaBP1 and the Nt*Eh*CaBP1 *EF*2 site as the model in this study since this set provided several advantages. 1) The sequence of the designed construct and that of the wild type were fairly similar so that relatively few single-site substitutions were required (Figure 1D). 2) We used an optimized set of protocols and availability of biophysical and structural data on the N-terminal domain construct of the protein. 3) The Nt*Eh*CaBP1 protein construct with two known Ca^2+^-binding sites allowed us to investigate the effect of one binding site on the other and hence to potentially understand the mechanism of cooperative binding in this protein. 4) The designed mutant sequence was unique.

**Figure 1.**
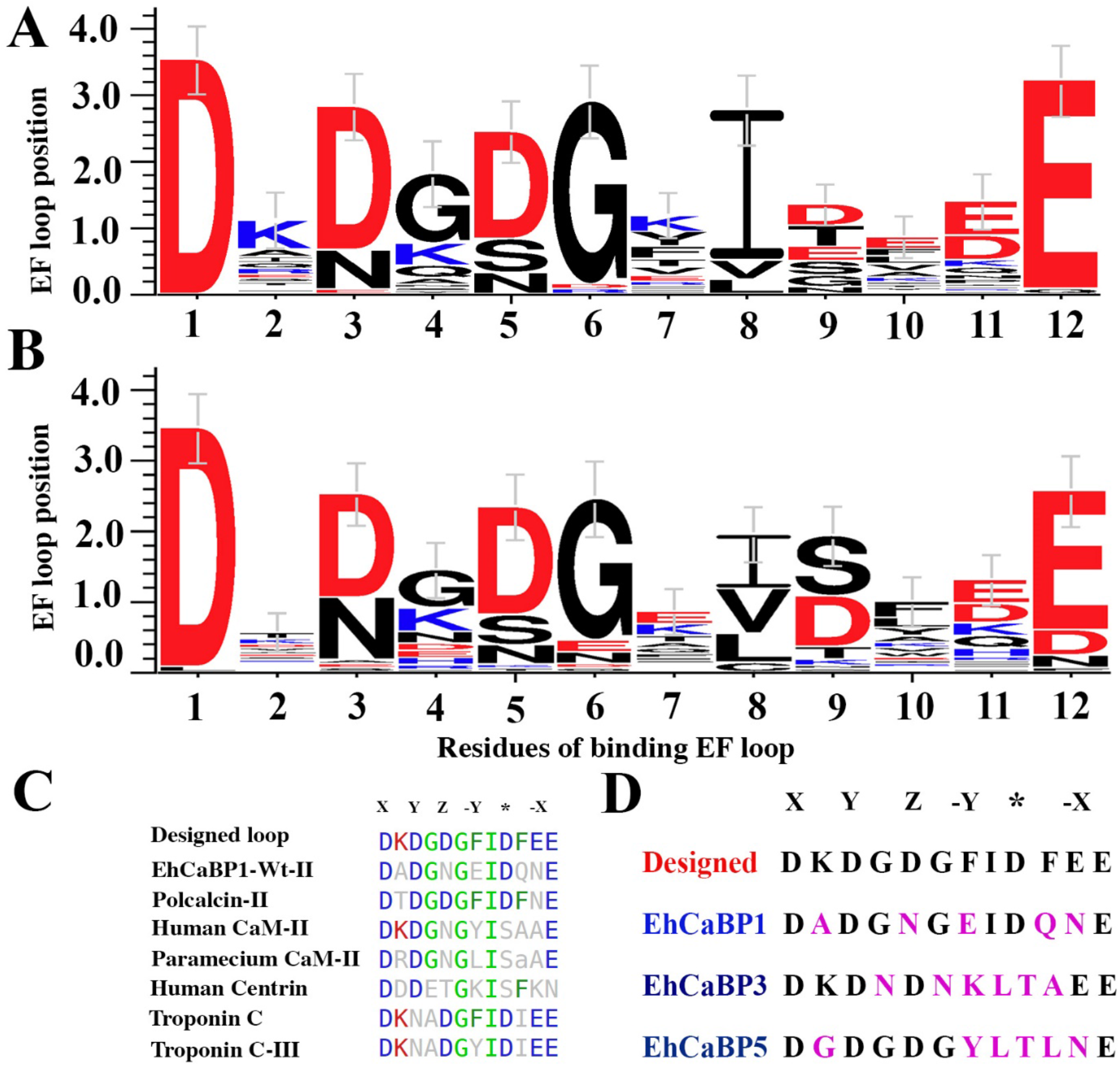
Sequence conservation and design of EF-hand loop sequence. A) Sequence logo of the residues forming high-affinity Ca^2+^-binding EF-hand loops from the published dataset with experimentally known binding affinities. B) Sequence logo of the residues forming of low-Ca^2+^ affinity EF-hand loops from the published dataset with experimentally known binding affinities[62]. The alignment showed certain position of the EF-hand loop to be relatively variable, but others to be quite conserved. C) The sequence alignment of the designed construct EF2 site with structurally closest homologs of EF-hand loops of NtEhCaBP1. D) The EF-hand loop sequences of the characterized and published constructs of EhCaBPs [31] [63][19] aligned with the designed peptide. This alignment showed the number of single-site substitutions (in pink) required to be performed in order to mutate the EF-hand loop to the designed construct.

### Cloning of the Nt*Eh*CaBP1 mutant

The gene fragments corresponding to the N-terminal domain of the EhCaBP1 protein were cloned by using an existing N-terminal clone of EhCaBP1 as a template in the bacterial expression vector pET28(b). The mutations were created (in NtEhCaBP1 EF-II) at positions 141, 150, 156, 165, and 168 by performing site-directed mutagenesis. The following primers were used for the mutation:

CaBP1 Mut K2 FP 5’-CAAATCTATTGATAAAGATGGAAATGG-3’

CaBP1 Mut K2 RP5’-CCATTTCCATCTTTATCAATAGATTTG-3’

CaBP1MutD5, F7 FP 5’CTATTGATAAAGATGGAGATGGATTTATTGATCAAAATGAATTTGC-3’

CaBP1MutD5, F7 RP 5’-GCAAATTCATTTTGATCAATAAATCCATCTCCATCTTTATCAATAG-3’

CaBP1Mut F_10_ FP 5’-ATTTATTGATTTTAATGAATTTGC-3’

CaBP1Mut F_10_ RP 5’-GCAAATTCATTAAAATCAATAAAT-3’

CaBP1Mut E_11_ FP 5’-ATTTATTGATTTTGAAGAATTTGC-3’

CaBP1Mut E_11_ RP 3’-GCAAATTCTTCAAAATCAATAAAT-3’

CaBP1 FP -5’-CATGCCATGGCAATGGCTGAAGCACTTTTTAAAG-3’

CaBP1 RP-5’-CGGCTCGAGGAGTGAAAACTCAAGGAATTCTTC-3’

*The* mutations were confirmed by DNA sequencing.

### Overexpression and purification of the NtEhCaBP1 EF-2 mutant

NtEhCaBP1 was expressed and purified as described in previously published literature [32]. The NtEhCaBP1 EF-2 mutant construct was transformed into *Escherichia coli* strain BL21 (DE3) for expression. Cells were grown in Luria-Bertani (LB) medium supplemented with 50 mg ml^-1^ kanamycin at 37°C. The culture was induced with 0.8 mM IPTG when the OD reached 0.7 at A_600_. It was then incubated at the same temperature for 3 h for further growth. Cells were harvested by subjecting by centrifugation at 7000 rpm for 10 min. The cell pellet was resuspended in suspension buffer (50 mM Tris pH 7.5, 2 mM EGTA). The cells were then lysed by freeze-thaw followed by sonication. A clear supernatant was obtained by carrying out centrifugation at 12000g for 30 min. The protein supernatant was passed through an ion-exchange chromatography using DEAE resin column, was pre-equilibrated with ten bed volumes of suspension buffer. The column was then washed with 30-40 ml of wash buffer (50 mM Tris pH 7.5, 5 mM NaCl) to remove nonspecifically bound proteins. Finally, the protein was eluted with elution buffer (50 mM Tris pH 7.5, 5 mM CaCl_2_). Further, an ion exchange purified protein was subjected to gel filtration chromatography using Superdex G-75 column pre-equilibrated in buffer containing, 50 mM Tris pH 7.5, 5 mM CaCl_2_. The Nt*Eh*CaBP1-EF2 mutant protein peak was eluted at 65.93 ml (Figure 3.1). The molecular weight was calculated for observed peak using standard protein marker of different molecular weights, supplied by Sigma Aldrich. The purity of the protein was checked using 15% SDS-PAGE.

### Isothermal titration calorimetry (ITC) to determine the Ca^2+^ binding affinity

Before performing ITC experiment both the wildtype (Wt-NtEhCaBP1) and mutant (Nt*Eh*CaBP1EF-2) proteins were prepared in Ca^2+^-free (apo form) form, as described previously [34] and the proteins were dialyzed against 10 mM of HEPES pH 7.4. ITC experiments were carried out using a Microcal ITC_200_ instrument from GE Healthcare. Experiments were performed at 30°C in 10 mM HEPES buffer (pH 7.4) at 1 mM of CaCl_2_ (titrant) concentration and 3.4 µM of the protein (analyte), either Wt-Nt*Eh*CaBP1 or the Nt*Eh*CaBP1EF-2 mutant were used. The volume of each titrant sample was 2 μl per injection and the fixed titrant pipette volume (40 µl) led us to perform 20 injections of titrant into a sample cell containing analyte (known concentration) as protein. The mixture was allowed to react for 2 min between injections and the heat release due to injection and dilution was obtained by titrating CaCl_2_ into buffer containing protein. A raw thermogram was generated with rate of change of heat with respect to time by using Microcal iTC200 software. The data were fitted using the Origin software Microcal Analysis Launcher supplied by Microcal. The plot of heat change with respect to molar ratio of [ligand]/[protein] was derived from the raw thermogram by using Origin software Microcal Analysis Launcher. Iterations were performed until the chi square value did not decrease further and was stable. The data were fit by sequential binding mode which gave number of binding sites (N=2) for ligand (Ca^+2^), association constant (K_a_), change in enthalpy (ΔH), and change in entropy (ΔS) values were obtained from the fitted data. Ligand to buffer titration was performed to check the background thermal noise of the buffer dilution. In order to decrease the thermal noise, a reference cell was filled with the same buffer used to prepare both the ligand and protein. The heat released and absorbed during titration was calculated by using the equation

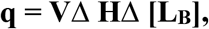

where q= change in heat resulting from the change in the concentration of bound ligand,

Δ[L_B_]= change in the concentration of bound ligand,

ΔH= change in enthalpy after ligand binding, and V= reaction volume.

### Crystallization of the NtEhCaBP1 EF-2 mutant

Prior to crystallization the SEC purified NtEhCaBP1 EF-2 mutant protein was concentrated to 15mg ml^-1^ by using a 3kDa-cut-off centricon filter. Crystallization trials were performed by using hanging-drop vapour-diffusion method in 24-well plates. 2µl of protein sample was mixed with an equal volume of precipitant solution in hanging drops and equilibrated against 500 ml of reservoir solution (precipitant). Initially the same crystallization condition was tried in which Wt-Nt*Eh*CaBP1 was crystallized [32], but it did not yield crystals. Rather the NtEhCaBP1 *EF*-2 mutant was crystallized in conditions similar to the crystallization condition of full length *Eh*CaBP1 [35]. The NtEhCaBP1 *EF*-2 mutant crystallize in the presence of MPD (58%-63%), 50mM sodium acetate (pH 5.0-5.5), and 5mM CaCl_2._

### X-ray diffraction, data collection, and processing and structure determination

NtEhCaBP1 EF-2 mutant crystals were soaked in cryoprotectant solution consisting of 65% MPD, 50 mM sodium acetate pH 5.3 and 5 mM CaCl_2_. Single crystals were picked up in cryoloop and flash-cooled in liquid nitrogen. Data were collected at ESRF DBT-BM14, France. The crystals diffracted to a resolution of 1.9 Å. Diffraction data were processed and scaled using HKL2000 [36]. Crystal belonged to the space group P2_1_2_1_2_1_, with unit cell parameters a= 44.6, b= 101.3, c= 107.4 Å. Matthews’s coefficient, V_M_, was 2.90 Å^3^Da^-1^, suggested presence of six molecules in the asymmetric unit, with a solvent content of 57.5% [37]. Structure was determined by molecular replacement method using Phaser program [38] and the assembled trimer of wild-type structure of *Eh*CaBP1 (PDB code 2NXQ) was used as a search model [35]. Overall, thirteen Ca^2+^ ions were identified in the electron density, two Ca^2+^ ions in each chain at the center of the EF-hand loop and one extra Ca^2+^ ion was observed which was not bound to an EF hand motif. All thirteen Ca^2+^ ions were included in the refinement. Structure was refined by REFMAC5 [39] and carrying out iterations of model building using the COOT graphics package [40]. For the final model, the *R*_work_ was 21.2 % and *R*_free_ was 24.9% and quality of structure using check using the program PROCHECK [41] showed it to have good stereochemistry with 97.6% of residues lying in the most-favored regions of the Ramachandran plot. Structure factor and refined model of NtEhCaBP1 EF-2 mutant were deposited in the Protein Data Bank [42] with the accession code 5XOP. Data collection and final refinement statistics are shown in Table 1.

**Table 1.** Crystallographic parameters and data collection statistics for the NtEhCaBP1 EF-2 mutant.

|  |  |
| --- | --- |
| X-ray source | ESRF BM14 Beamline |
| Wavelength (Å) | 0.9788 |
| Space Group | P2<sub>1</sub>2<sub>1</sub>2<sub>1</sub> |
| Cell parameters (Å, °) | a = 44.6, b = 101.3, c = 107.4<br>$\alpha = 90, \beta = 90, \gamma = 90$ |
| No. of molecules in an asymmetric unit | 6 |
| Resolution range (Å) | 50-1.9 (1.93-1.9) |
| Total Reflections | 301767 (15064) |
| Unique Reflections | 39381(1966) |
| Completeness (%)<sup>a,b</sup> | 99.9 (100) |
| Redundancy | 7.7 (7.7) |
| Mean I/$\sigma$ | 33.0 (1.9) |
| Rmerge (%)<sup>b,c</sup> | 7.5 (88.8) |
| **Refinement** |  |
| R<sub>work</sub> (%) | 21.2 (27.1) |
| R<sub>free</sub> (%) | 24.9 (33.6) |
| Mean B-factor | 42.5 |
| No. of atoms<br>Protein/Ca/water/MPD | 3128/13/174/1 |
| Bond length (Å) | 0.015 |
| Bond angles (°) | 1.71 |
| Dihedral angle (°) | 17.6 |
| **Ramachandran plot statistics** |  |
| Favoured region (%) | 95.5 |
| Additional allowed region (%) | 4.2 |
| Generously allowed regions (%) | 0.3 |
| Outliers | 0.0 |
| **PDB code** | **5XOP** |

### Structure and sequence analyses

We performed the sequence alignments using Clustal Omega and BioEdit [43, 44]. The structural alignment was performed using the Dali server [45]. Protein-protein interactions between the subunits were calculated using the *PDBsum* webserver [46] and Dimplot [47]. The Ca^2+^ coordination distances and angles were calculated using LIGPLOT [47] and the PLIP server [48] as well as using PyMol software [49]. Representations of the structure were prepared by using PyMol, Chimera [50] [45] and Photoshop software [51]. The individual residues involved in interactions of both the structures were analyzed by using the Arpeggio web server [52] and Hydrogen Bonds Computing Server (HBCS) [53]. The contact maps from the protein structures were calculated by using the Residue Interaction Network Generator (RING) web server [54].

### Cross-correlation and dynamic network analysis

Network and dynamics cross-correlation map (DCCM) analyses were performed by using the Bio3D package [55]. Here, the proteins network systems were used to show insights into the structural relationships between three EF-hand-containing protein systems exhibiting different binding affinities. The dynamic networks were generated by using an ensemble of structures showing cross correlation between intramolecularly connected residues. We used normal mode analysis (NMA) [25] to obtain insights into the coupled motions and allosteric effects directly influencing the cooperativity of the two Ca^2+^-binding sites in each protein. Furthermore, the cross-correlation matrix was calculated from the NMA, and a DCCM [55] was generated for three proteins using the Hessian matrix. C_ij_ values of the matrix close to 1 means the fluctuations of residues i and j are completely correlated (same period and same phase), a C_ij_ close to -1 suggests that the fluctuations of residues i and j are completely anti-correlated (same period and opposite phase), and a C_ij_ value equal to 0 indicates the fluctuations of i and j are not correlated. In the figure made, C_ij_ values are indicated in the right panel, and the cyan color signifies the positive correlation and pink shows the negative or anti correlation. Such a depiction allows for the identification of highly intramolecularly connected residues clustered as individual communities shown in the same color. The community clustering (clusters as communities in a graph), which deals with the graph partitioning problem, has been solved by the Girvan-Newman edge-betweenness approach which is implemented in Bio3D to generate the network [55]. The residues belonging to the communities/clusters in the graph are also shown in different color codes along with position of each residue in the DCCM on the X axis and Y axis.

## Results

### Roles of EF-hand loop residues in Ca^2+^ binding

Mutations occurring in the sequences of many critical genes form the major driving mechanism leading to the evolution of complex organisms. These mutations have been seen in different protein families across all biological systems. But mutations are not prevalent in certain locations of proteins such as their structural cores, and the conserved nature of these residues are suggestive of their important functional roles [56]. Enhancing the ability of a protein to carry out a specific function requires an understanding of the functional domains and the roles of conserved residues. The EF-hand is a well-studied motif having many conserved residues, where the conservation is particularly strict at the Ca^2+^-binding site. The well-studied high preference of certain amino acid residues at particular positions of the motif, such as aspartate at the beginning of the motif and glutamate at the end indicates that some ligands are indispensable for Ca^2+^ binding [17, 57]. These negatively charged residues placed at many positions in a short stretch of sequence makes an ideal site for the binding of a positively charged divalent ion such as Ca^2+^ or magnesium. In physiological conditions, both of these ions are present in the cell in high concentrations, yet the selectivity for the metal ion is highly specific in EF-hand proteins [58, 59]. Ca^2+^-binding loops have evolved to reflect biological functions that require of the proteins a range of binding affinities [5]. In order to understand the role of individual residues in the Ca^2+^-binding loop, we classified our data into two groups, namely high-affinity sites (HASs) and low-affinity sites (LASs) and aligned the sequences within each group to derive sequence motifs of the two groups (Figure 1). Figures 1A and 1B illustrate, for the two groups, the distributions of residues at each position of the EF-hand loop of the Ca^2+^-binding site. The first position (X) of the loop is dominated by aspartic acid (D), known to be indispensable for Ca^2+^ binding due to its being involved in the critical hydrogen-bonding network and due to its providing an oxygen for coordinating the Ca^2+^ [60]. Some of the EF-hand Ca^2+^-binding sites with a non-aspartate residue at the X position have shown low or even no affinity for Ca^2+^. This observation has also been seen for numerous mutational studies of different EF-hand proteins [60]. Comparison of the high-affinity and low-affinity sequences (figure 1A, 1B) suggests that positively charged lysine in the second position (non-interacting site) of the loop is present more often in HAS sequences than in LAS sequences. In this position of the LAS sequences showing the occurrence of a variety of amino acid residues such as alanine and isoleucine. In the third position (Y position), aspartic acid occurs most often followed by asparagine. The eighth position is mostly occupied by the hydrophobic residue isoleucine, valine and leucine. A close inspection of the sequences indicates a relatively high occurrence of serine at the 9^th^ position of LAS sequences but not HAS sequences. In the twelfth position, glutamic acid is conserved in HAS sequences, not surprising since this residue provides a bidentate side-chain ligand for Ca^2+^ binding. In a few LAS sequences, aspartic acid is present in the twelfth position. Aspartic acid at the 12^th^ position sometimes requires, because of the short side chain, one more water molecule to coordinate Ca^2+^, resulting in a lower affinity for the Ca^2+^ due to increased solvent entropy [60]. The analysis suggested that Ca^2+^-binding loop residues that do not directly coordinate the Ca^2+^ nevertheless have important roles for the binding of Ca^2^, including stabilizing the loop, contributing to the extensive intricate hydrogen-bond network required for the proper folding of the loop, and properly positioning the negatively charged residues of the loop into the three-dimensional arrangement best suited for Ca^2+^ binding [60].

### Scoring scheme to assist modeling of the EF-hand Ca^2+^ binding site

In an earlier study, we were able to classify EF-hand domains based on their affinities (high, low and none) for Ca^2+^ [61]. This classification was done on the basis of datasets trained in machine learning, specifically based on support vector machines (SVMs). The correlations derived from the use of various amino acid indexes showed high prediction accuracies when tested against many *Eh*CaBPs [62]. In the present study, we extracted the margin distance from the decision boundary for each prediction, generated PSSM values, and incorporated the SVM_Mar_ and the PSM_LogL_ scores as added features to the previously developed method. Furthermore, we calculated the margin distances and conservation scores for all of the sequences of the three different previously published datasets consisting of 121 unique Ca^2+^-binding EF-hand loops. The details of the predictions for each site along with the new scores are shown in Appendix I. The scores indicated the possibility of classifying individual sites in an EF-hand protein on the basis of their distance from the decision boundary. The conservation score shows the probabilities of the residues occurring frequency.

### Designing a high binding affinity EF-hand loop

So far, several mutational and design studies on EF-hand proteins have yielded distinctive impacts on their range of binding affinities, thus influencing the functions of those proteins [19, 23] [59]. In many of these studies, mutations were made to help assess the Ca^2+^-binding capability of the protein. Substitutions in the first (X) and last (-X) positions of the binding site did yield desirable results, as these ligand positions are extremely important and require negatively charged chelating residues for proper function [57, 63]. In order to validate our prediction method, we designed a unique EF-hand loop, that is not presents in any protein sequence database (Figure 1C). The sequence design was executed by substituting every amino acid residue in each of the 12 positions of EF-hand motif. Each mutant was analyzed, and only those that increased the score were taken to the next iteration (see diagram for schematic representation). The designed binding loop with the amino acid residue sequence 1-DKDGDGFIDFEE-12 showed a high SVM_Mar_ score of 2.694 and PSM_LogL_ score of 6.46, “, both comparable to those of the Ca^2+^-binding sites with high affinity from the literature [23]. Sequence alignment performed with EF-hand containing proteins from different families (Figure 1C) suggested that the designed loop is most similar to the Ca^2+^-binding pollen allergen [64], and similar to the structure of the WtNt*Eh*CaBP1 protein, which shows a head-to-tail arrangement with domain-swapped EF-hand pairing. We inserted the EF-hand loop sequence having the high predicted Ca^2+^ binding affinity into the well-characterized WtNt*Eh*CaBP1 Ca^2+^-binding site to attempt to manipulate the Ca^2+^ binding affinity of this protein. We specifically mutated the sequence of the 2nd EF-hand motif of the Ca^2+^-binding loop of Nt*Eh*CaBp1. The Nt*Eh*CaBP1-EF2 loop was selected because of the greater similarity between its sequence and that of the designed loop than between the sequences of the loops of other characterized Ca^2+^-binding proteins of *E. histolytica* and that of the designed loop (Figure 1D). The crystal structure of the N-terminal construct of Nt*Eh*CaBP1 has two Ca^2+^-binding sites, while the full-length protein has four Ca^2+^-binding EF-hand motifs [32, 35]. In a previous study, ITC experiments using full-length protein showed four binding sites with two sites displaying high Ca^2+^-binding affinity and two sites with low binding affinity [65]. Calculations performed on the first Ca^2+^-binding motif, that is the EF-1 (1st EF-loop-DVNGDGAVSYEE), yielded SVM_Mar_ and PSM_LogL_ scores of *-1*.*045* and *4*.*976* and hence predicted EF-1 to have a relatively low binding affinity for Ca^2+^. Calculations performed on the 2^nd^ EF-loop, i.e., with the sequence DADGNGEIDQNE, yielded SVM_Mar_ and PSM_LogL_ scores of 1.001, and hence predicted the 2^nd^ EF-loop to have a Ca^2+^-binding affinity lower than that of the first, i.e., EF-1, loop. To preserve the proper folding of the molecule the construct was built with two EF-hand loops. We were also interested in understanding how synthetic EF-hand loops influence cooperative binding in this system. In order to accomplish these aims and to understand the biophysical characteristics of the protein containing the engineered loop, we purified both WtNt*Eh*CaBP1 and the Nt*Eh*CaBP1 mutant (EF2) and compared them by performing biophysical and computational experiments.

### NtEhCaBP1-EF2 mutant binds Ca^2+^ with higher affinity

To determine whether incorporating mutation in EF-2 loop of Nt-*Eh*CaBP1 (Nt-EhCaBP1-EF2 mutant) enhanced the Ca^2+^-binding affinity, we performed the ITC experiments. Initially we performed the titration in presence of 1mM of CaCl_2,_ but WtNt-EhCaBP1 did not shows the Ca^2+^ binding (data not shown) and curve did not achieve the saturation. In contrast, the binding of Ca^2+^ to Nt-*Eh*CaBP1-EF2 mutant is strong, and a well saturated curve obtained (Figure2B). To get better binding curve for WtNt-EhCaBP1, titration was performed with increasing concentration of Ca^2+^ and titration at 25mM of Ca^2+^ gave a better binding curve, with very low association constant than Nt-EhCaBP1-EF2 mutant form of protein (Table 2). The low association constant suggest that binding affinity is very weak in case of WtNt-EhCaBP1.

**Figure 2.**
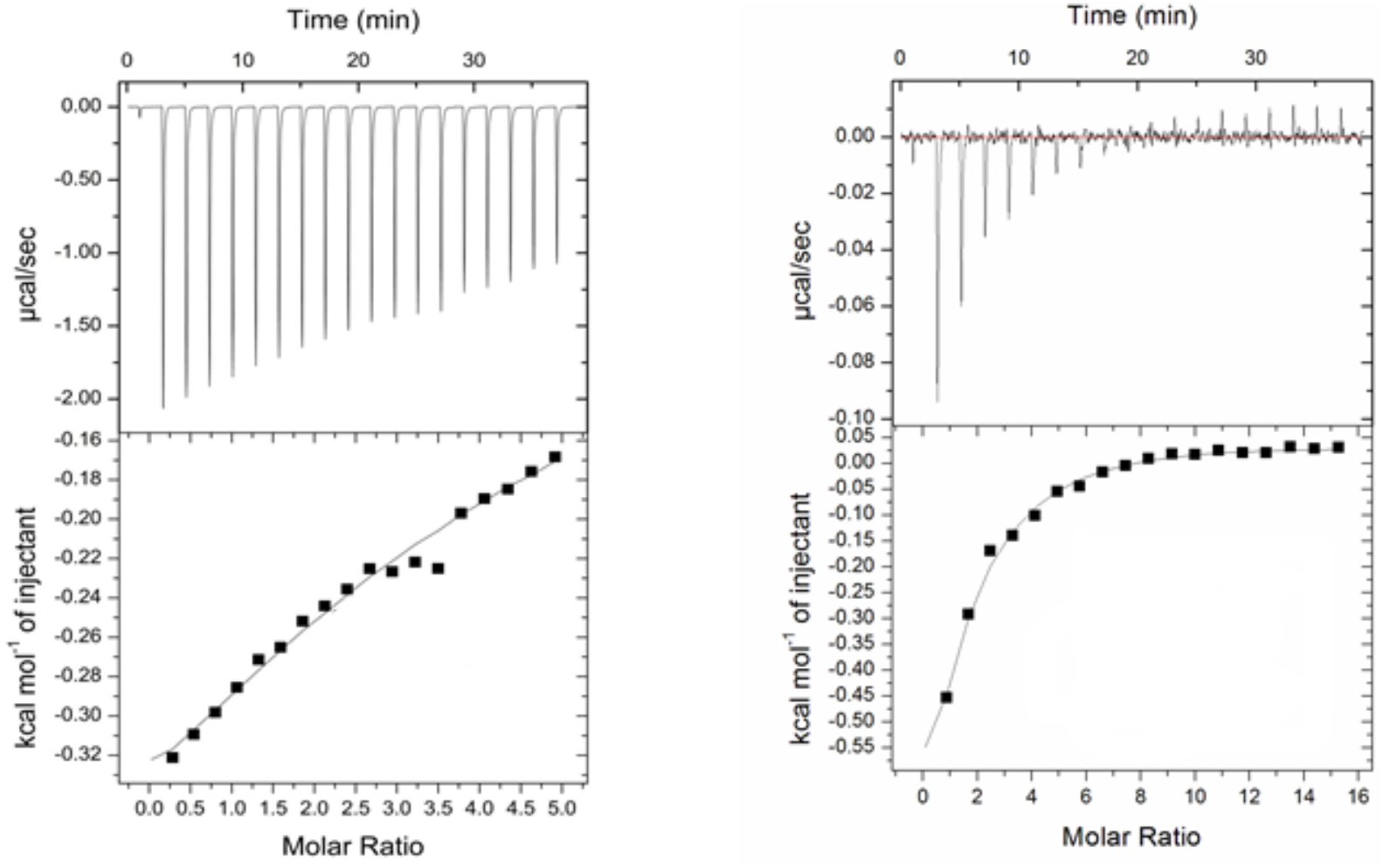
Isothermal titration calorimetric analysis of the binding of Ca^2+^ to the A) apo Nt*Eh*CaBP1 and B) apo Nt*Eh*CaBP1 EF-2 mutant. Plots are shown of kcal of heat absorbed/released per mole of injected CaCl_2_ at 30°C. For all titrations, the top panels represent the raw data (power versus time) and the bottom panels represent integrated binding isotherms. The solid line in each bottom panel represents the best nonlinear fit to the experimental data. The thermodynamic parameters obtained from these plots are summarized in Table 1.

**Table 2.** The hydrogen bonds formed between the trimer1 (Chain: A, B, C) and trimer2 (Chain: D, E, F).

| Trimer 1 |  | Trimer 2 |  | Distances |
| --- | --- | --- | --- | --- |
| **GLN-36 (NE2)** | Chain A | Ile-65 (O) | Chain D | 3.06 |
| **GLN-36 (NE2)** | Chain B | Ser-64 (O) | Chain F | 3.36 |
| **GLN-36 (NE2)** | Chain C | Ile-65 (O) | Chain E | 3.28 |

The binding constant of Nt-EhCaBP1 EF-2 mutant is 45 and 77-fold higher than the WtNt-EhCaBP1. The WtNt-EhCaBP1 curve shows the exothermic process with a favorable change in enthalpy for site-1 (ΔH1, -760.4) and unfavorable change in enthalpy (*ΔH*2, 4857 kcal/mol) for site-2 (Table 1). The entropy at site-2 is very low (−7.38) which suggest the binding of Ca2+ at site-2 is not bringing any large structural or conformational changes.

The Nt-EhCaBP1 EF-2 mutant curve shows an exothermic process for site-1 with a favorable change in enthalpy (ΔH1, -1866) whereas site-2 represent an endothermic process with unfavorable change in enthalpy (*ΔH*2, 5699 kcal/mol) and comparatively higher entropy than the site-1 (Table 1). The unfavorable enthalpy and higher entropy indicate that Ca^2+^ binding at site 2 undergoes large conformational change as compared to site 1. Usually, the Ca^2+^ binding protein shows the cooperativity when their EF-hands bind with Ca^2+^ ion i.e., binding of metal ion at one site induced the binding at second site. As we observed that NtEhCaBP1 is unable to bind Ca^2+^ or has weak binding affinity but in case of Nt-EhCaBP1-EF2, mutation induced the binding of Ca^2+^, therefore a µM range binding affinity obtained at both sites. This difference in binding affinity in both form of protein suggests that Nt-EhCaBP1 EF-2 mutant shows the phenomenon of cooperativity, which leads to obtain a better binding curve in comparison with Nt-EhCaBP1.

**Table 1.**
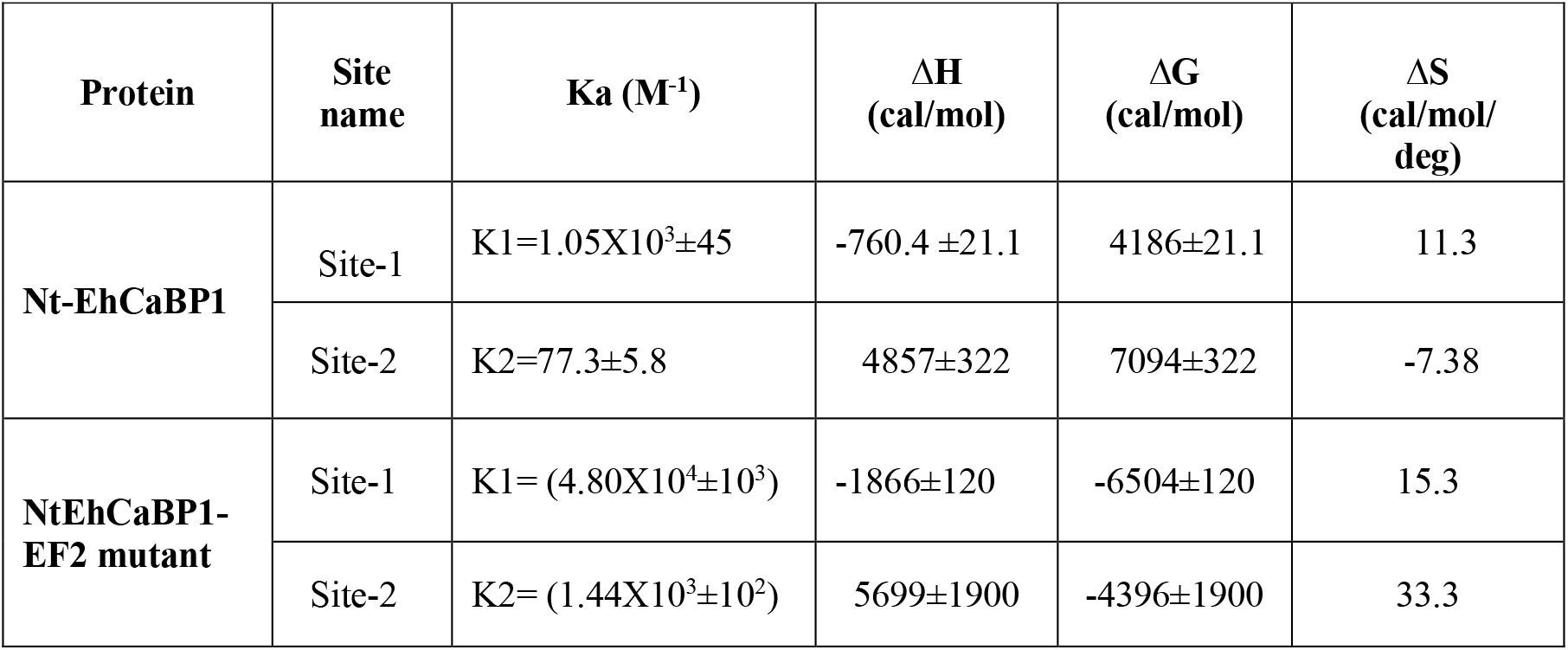
Summary of binding constants and thermodynamic parameters obtained from the Ca^2+^-binding isotherms of ITC studies of Nt*Eh*CaBP1 and Nt-*Eh*CaBP1 EF-2 mutant at 25°C.

### Solution state oligomerization and three-dimensional structure of NtEhCaBP1 EF-2 mutant

In our previous reports it has been shown that either full length EhCaBP1 or its C-terminal deleted mutant (NtEhCaBP1) form of protein always forms a trimer in solution or in crystal form [32, 35]. In contrast to our previous observation, we found that while purifying Nt*Eh*CaBP1 EF-2 mutant protein differed in oligomeric state with respect to Nt-EhCaBP1. The size exclusion profile (SEC) of Nt*Eh*CaBP1 EF-2 mutant protein shows a peak correspond to molecular mass of about 42-44 kilodalton (kDa). As, the molecular wight of NtEhCaBP1 is 7.0 kDa, the SEC profile suggest the formation of hexamer in solution (Figure 3A).

**Figure 3.**
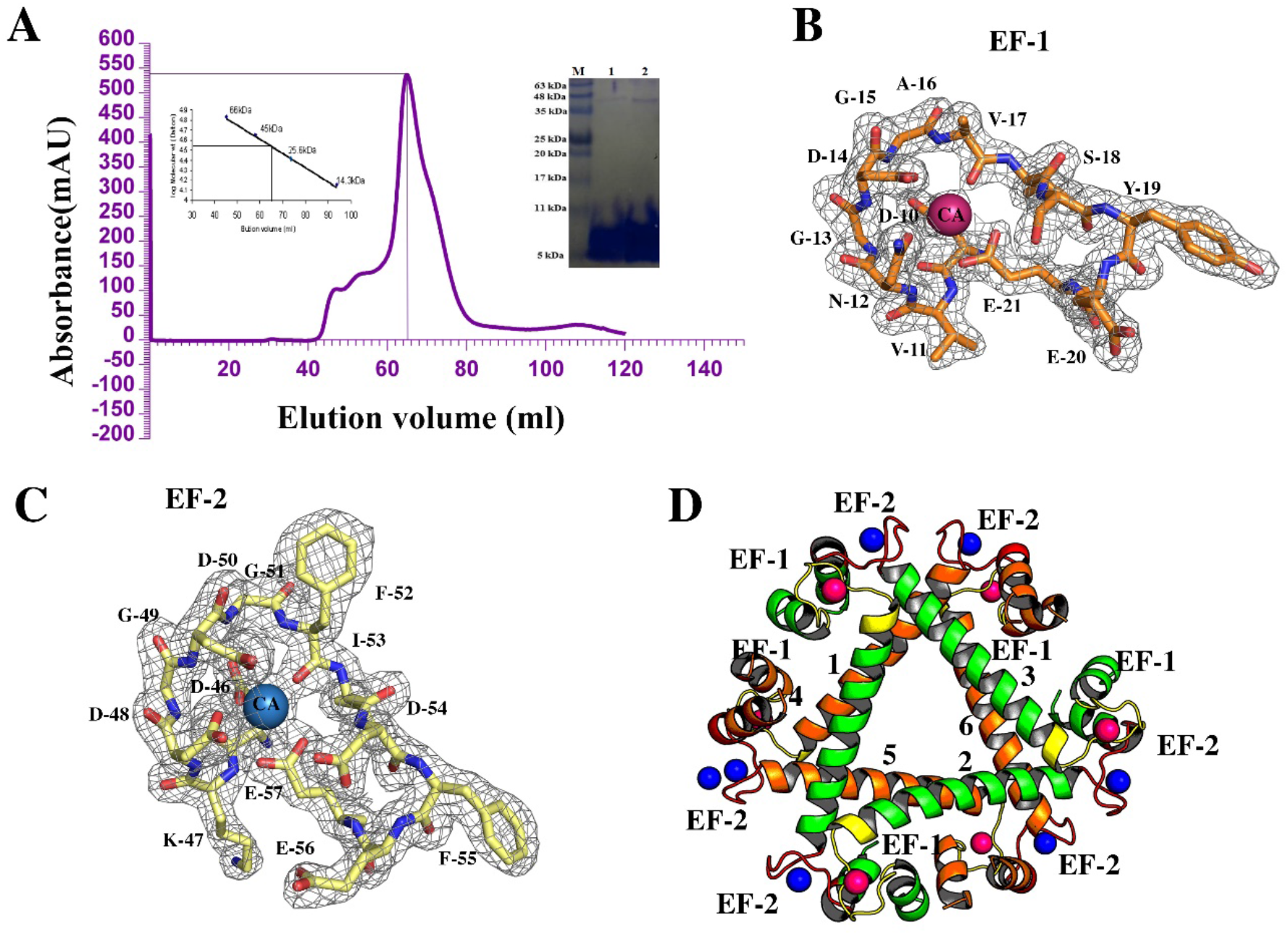
A) Size exclusion chromatography (SEC) profile of the Nt-*Eh*CaBP1-EF2 mutant) The SEC peak of Nt-EhCaBP1-EF2 mutant obtained from Superdex G-75 columns, shows the peak at elution volume of 64 mL, which corresponding to molecular weight of around 39kDa. Considering the molecular weight of NtEhCaBP1-EF2 monomer is approximately 7kDa, the eluted peak at 63ml indicates the formation of hexamer in solution. B, C) Electron density maps and structural models of the EF-1 and EF-2 Ca^2+^-binding loops of Nt*Eh*CaBP1 EF-2 mutant with bound Ca^2+^. The images specifically show the portions of the 2Fo-Fc electron density maps, each at a 1.0 σ cutoff, corresponding to the residues of the N-terminal (EF1-orange) and C-terminal (EF2-yellow) Ca^2+^-binding loops. All twelve amino acid residues (labeled) comprising each binding loop are represented in stick form inside each electron density map. D) The structure of the Nt*Eh*CaBP1-EF2 mutant forms hexamer, assembled via two trimers.

Consistent with the solution state hexameric form, crystal structure also shows presence of six molecule in the asymmetric unit, and it is arranged in the form of two trimer, which leads to hexamer formation. The structure is well refined, and a representative electron density map of EF-1 and EF-2 are shown in (Figure 3B & 3C). Structural analysis suggested that each trimer in Nt-EhCaBP1 EF-2 mutant structure adopted similar trimeric arrangement as formed in case of Wt-NtEhCaBP1 structure [32] (Figure 3D).

In the hexameric state, one trimer (interface-A) of the Nt*Eh*CaBP1-EF2 mutant interact with the other trimer (interface-B) (Figure 4D) where chain A was observed to interact with chain D (Figure S1) in the same manner chain B interact with chain F and chain C with chain E. The residues present on the binding interface of both the trimers forms multiple hydrogen bonds and non-bonded interactions. The oxygen atom from the C-terminal residues Ile-65, Ser-64 and Ile-65 are involved in hydrogen-bond interactions (Table 2) with nitrogen atom of glutamate-36 of the central helix (Figure 4E). This interaction brings the both the trimer in close proximity thereby leading to hexamer formation, in solution as well as in crystal form.

**Figure 4.**
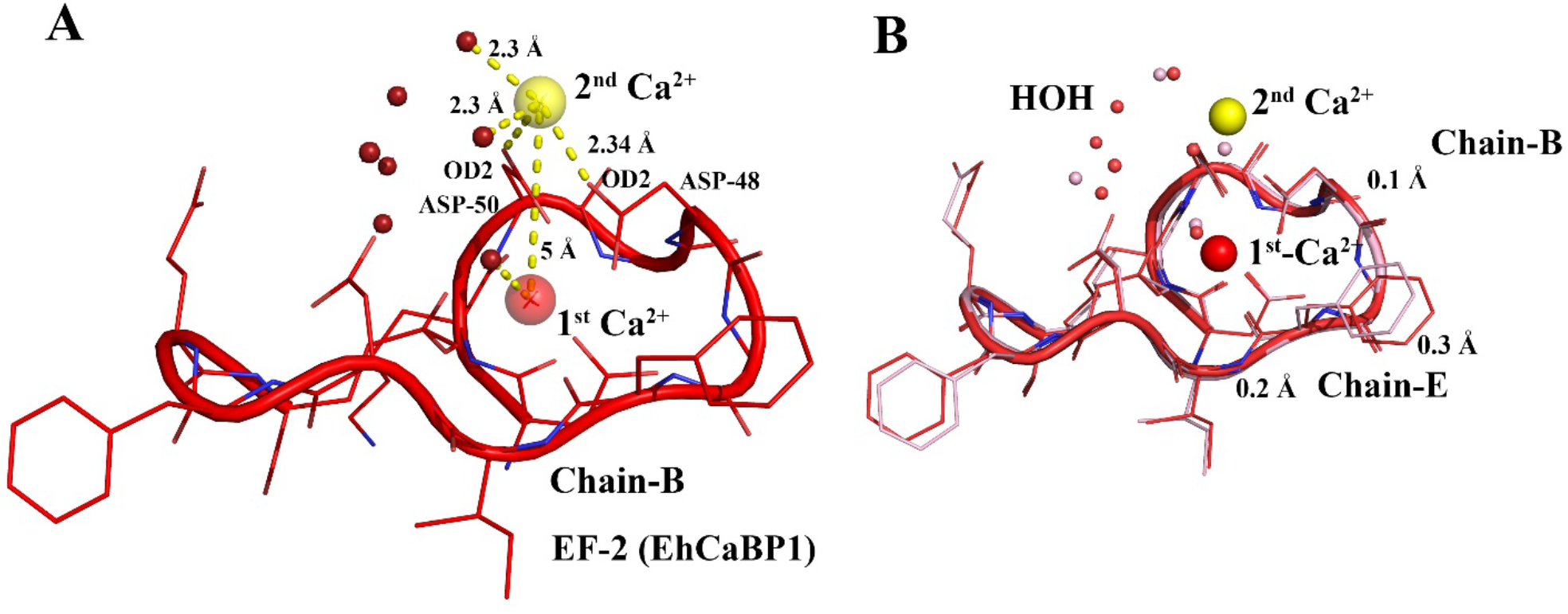
Two Ca2+ ions bound to the designed EF-2 of chain B of the Nt*Eh*CaBP1-EF2 mutant. A) The modified EF-2 loop of chain B of the NtEhCaBP1-mutant structure shown in a cartoon and lines representation and showing the locations of the two bound Ca2+ ions. The 2^nd^ Ca^2+^ ion positioned outside of the Ca^2+^-binding coordination sphere (colored in yellow). Both Ca^2+^ ions are separated with a distance of 5Å, 2^nd^ Ca^2+^ ion interact with oxygen atom of Asp48, Asp50 and two water molecules, both are located at a distance of 2.30 Å from Ca^2+^ ion.

The interactions between the three assembled domains in each trimer of Nt-EhCaBP1-EF2 mutant structure were found to be similar as it was observed in case of Wt-Nt*Eh*CaBP1 structure. There are some additional hydrogen bonding interactions were found in hexamer of NtEhCaBP1-EF2 mutant structure. The various interactions involved in hexameric state of Nt-EhCaBP1-EF2 structure are shown in supplementary figures S1 and S2 (Table S4 -S5).

### An extra Ca^2+^ ion bound outside of EF-hand motif

Apart from the usual Ca^2+^ ions occupied at each EF-hand motifs, an extra Ca^2+^ ion was observed at the 2^nd^ EF-hand motif of chain B. This extra ion occupied space outside of the regular Ca^2+^-binding coordination sphere. In order to understand the binding of two Ca^2+^ ions to one EF-hand motif, we calculated and plotted all the atomic interactions involving each of the two Ca^2+^ ions (Figure 4A). The first Ca^2+^ ion was observed to interact with X, Y, Z, -Y, -Z, and -X, including specifically with one of the side-chain oxygens (OD1) of each of the aspartate D-48 (Y) and D-50 (Z), to form a pentagonal bipyramidal geometry like conventional EF-hand loop. Whereas the second Ca^2+^ ion is bound outside of EF-hand motif and interacts with the side-chain oxygen atom (OD2) of D-48, D-50 and two water molecules. Most likely these two water molecules stabilize this extra Ca^2+^ ion (Figure 4A).

We also compared the EF2 loop of chain B with the EF2 loop of chain E (Figure 4B) including all the well-ordered water molecules within 5 Å of each loop. This analysis clearly indicated that the binding of the extra calcium to the EF2 loop of chain B was associated within the proximity of two water molecules to this calcium. In chain B the two water molecules interacted with the extra calcium at typical distances 2.30 Å. The occurrence of clusters of water or water-mediated hydrogen-bonding interactions have been seen in a greater percentage of non-EF-hand calcium-binding proteins (33.1%) than EF hand-containing proteins (13.1%). Our current comparison of the EF-2 loops of the different chains of the Nt*Eh*CaBP1-EF2 mutant suggested that binding of the additional calcium in chain B was not due to any additional structural changes but due to two close water-mediated interactions with the Ca^2+^ ion stabilizing the binding of Ca^2+^ outside of the EF-hand motif.

### Mutation induced bend in the third helix (H3) of NtEhCaBP1 EF-2 structure

To investigate whether the mutation induced any structural changes in the overall structure or in particular EF-hand motif, we superimposed the NtEhCaBP1 EF-2 mutant structure on Wt-NtEhCaBP1 by using PyMol. An alignment of all 66 Cα atoms yielded an RMSD of 0.96 Å. Structural superimposition shows the difference in the orientation of the third α-helix (i.e., H3) (Figure 5A). Distance measurement between second helix and the C-terminal part of third helix indicates that Nt*Eh*CaBP1 EF-2 mutant structure acquire 76.3 Å whereas the wildtype structure shows the distance of 83.4 Å, suggesting the occurrence of a distinctive structural change due to this mutation of the Ca^2+^-binding sites. We calculated this difference in the orientation of the third helix by measuring the angle between CD2 atom of residue Leu-36, CD1 atom of residue Ile-44, and the OE-1 atom of residue Gln-65 (Figures 5B & 5C). This measurement suggests a difference of about 7.1°, between the orientation of the third helix with respect to helix-2, in the mutant structure (Figure 5C).

**Figure 5.**
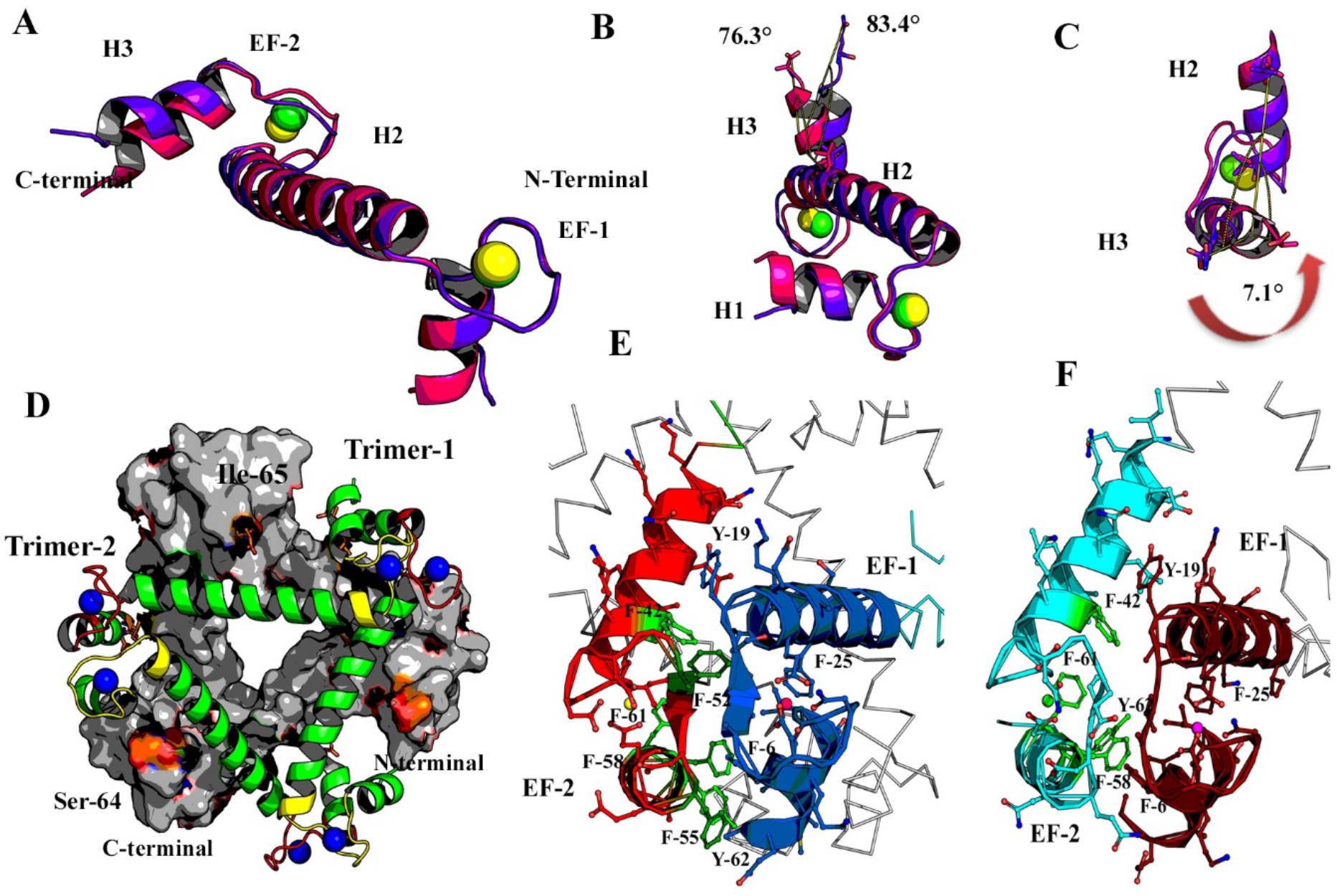
Superimposition of the WtNt*Eh*CaBP1 and Nt*Eh*CaBP1 mutant structures. A) The structure of WtNt*Eh*CaBP1 is shown in violet and that of the Nt*Eh*CaBP1 mutant in pink. B) & C) Two additional views of this superposition, with these views showing highlighting the different angles between the C-terminal helix and central helix for the two proteins. The change of the orientation of the C-terminal helix is shown with a curved red arrow. D) Trimer 1 (chains A, B, C) is shown in surface representation and trimer 2 (D, E, F) is shown in cartoon representation forming hexamer. The c-terminal residues are shown at the binding interface of protein-protein interactions. E) The interactions of the EF-1 and EF-2 motifs from two subunits of the Nt*Eh*CaBP1 mutant were found to mimic the pairing required for the proper functioning of the EF-hand proteins [53]. The hydrophobic residues are labeled and the residues in the EF-2 loop are shown in green. F) Similarly, the two chains from the trimer of NtEhCaBP1 is shown with residues forming the hydrophobic core.

### Hydrophobic interactions leading to positive cooperativity

Hydrophobic interactions are one of the driving forces in EF hand proteins due to their role in the cooperative binding of the two EF hand motifs to each other [13, 17, 21]. The locations of hydrophobic residues in the Nt*Eh*CaBP1 structure and in the Nt*Eh*CaBP1 mutant structure interacting are highlighted in green (Figure 5E and 5F), where the four-helices forming the hydrophobic core are tightly associated with each other. These interactions mostly involved phenylalanine and tyrosine residues, which appeared to play an important role in stabilizing the EF-hand loop by forming interactions between one EF hand loop and the other (Figure 5E and 5F). The interactions were aided by the formation of an antiparallel EFβ-scaffold (Figure 5E), which helps the loop to close while folding without causing large structural change[64]. The stronger allosteric interactions between the odd-numbered and even-numbered EF loops were found to lead to a greater calcium binding affinity; in the Nt*Eh*CaBP1 mutant, these interactions appeared to be further enhanced due to the presence of two additional aromatic residues (E52F and Q55F). The observation of conserved hydrophobic residues at the 6^th^ and 22^nd^ positions of EF hand motifs suggested the communication between the two EF hands to be chiefly mediated by the noncovalent interactions.

### Changes in overall charge distribution

To understand the differences between the charge distribution in the mutated loop and that in the wildtype loop, we calculated the overall surface charge distribution of the Nt*Eh*CaBP1 mutant and that of WtNt*Eh*CaBP1 (Supplementary Figure S3) using ABPS plugin in PyMol. The analysis indicated a similar charge distribution in the calcium-binding loop of the mutant as in that of WtNt*Eh*CaBP1, except for the replacement of wildtype negatively charged glutamate at position 52 with hydrophobic phenylalanine. We calculated the number of oxygen atoms available to the mutant and wild type EF-2 and found more oxygen atoms in the wild type than in the mutant protein (Figure S3). This result suggested that increasing the binding affinity for Ca^2+^ does not necessarily depend upon introducing more negatively charged atoms but in creating an optimum distribution of charges where the stabilizing hydrophobic interactions overcome the destabilizing electrostatic interactions caused by the negatively charged residues in close proximity [66].

### Comparison of EF-hand-2 of Nt-EhCaBP1-EF2 mutant structure with Wt-NtEhCaBP1

To understand if enhanced Ca^2+^ binding observed in ITC data of Nt-EhCaBP1-EF2 mutant protein, contributed any changes in Ca^2+^ coordination sphere we compared the distances and angles of the bound Ca^2+^ with their respective interacting residues of the EF-2 loop of the mutant protein with the EF-2 of Wt-Nt*Eh*CaBP1(PDB code 2NXQ). The Ca^2+^ coordination distance (d1) acquired by various residues in Nt-EhCaBP1-EF2 mutant structure were comparatively small with respect to the Ca^2+^ coordinating distance (d2) acquired by respective residues of Wt-Nt*Eh*CaBP1 structure (Table 3). Compared to Wt-Nt*Eh*CaBP1 structure, each Ca^2+^ coordinating residues of Nt-EhCaBP1-EF2 mutant structure shows the difference of ∼0.1Å-0.5Å, suggesting that reduced distance in Ca^2+^ of Nt-EhCaBP1-EF2 structure attribute the high Ca^2+^ binding affinity.

**Table 3.** List of the Ca^2+^-binding ligands at the conserved positions and water molecules of the Nt*Eh*CaBP1-mutant EF-2 loop and WtNt*Eh*CaBP1 EF2-loop as shown in the figure 5 (A and B) along with the distances (Å) between the interacting atoms/ residues and the angles (Θ) with respect to the water molecule. Specifically listed are residues of EF-2 of chain A interacting with Ca^2+^ through their coordinating side-chain oxygens.

| Residue | Position | Angle (Θ) | AA | Distance (d1) | Angle (Θ1) | AA | Distance (d2) | Angle (Θ2) |
| --- | --- | --- | --- | --- | --- | --- | --- | --- |
| Binding Site (EF-2) |  |  | NtEhCaBP1-mutant |  |  | WtNtEhCaBP1 |  |  |
| 46 | **X** | OD1- HOH-CA | ASP | 2.34 | 172.6 | ASP | 2.64 | 158 |
| 48 | **Y** | OD1-HOH-CA | ASP | 2.4 | 95.7 | ASP | 2.84 | 102.4 |
| 50 | **Z** | OD1-HOH-CA | **ASP** | 2.22 | 84.7 | **ASN** | 2.70 | 73.6 |
| 52 | **-Y** | O-HOH-CA | **PHE** | 2.27 | 97 | **GLU** | 2.56 | 105.3 |
| 57 | **-Z** | OE1-HOH-CA | GLU | 2.66 | 95.6 | GLU | 2.7 | 109.1 |
| 57 |  | OE2-HOH-CA | GLU | 2.5 | 79.5 | GLU | 2.4 | 115.2 |
| water | **-X** | NA | HOH | 2.32 | NA | HOH | 2.92 | NA |

Also, the oxygen-water-Ca^2+^ angles (Θ1), of Nt-EhCaBP1-EF2 mutant structure shows the difference, for example Asp-46, interacting from the top of the pyramidal geometry showed an HOH-CA-OD1 angle of 172.6° compared to 158° for Nt*Eh*CaBP1-EF2 (Figure 6 A & 6B). The other coordinating contacts showed a variation of ∼15° of angles (80° to 95°) amongst themselves (Table 3). These differences in bond lengths and angles between the mutant and wildtype structures may have caused the difference in ITC results, where the Nt*Eh*CaBP1-EF2 mutant protein sites showed higher binding affinities for Ca^2+^.

**Figure 6.**
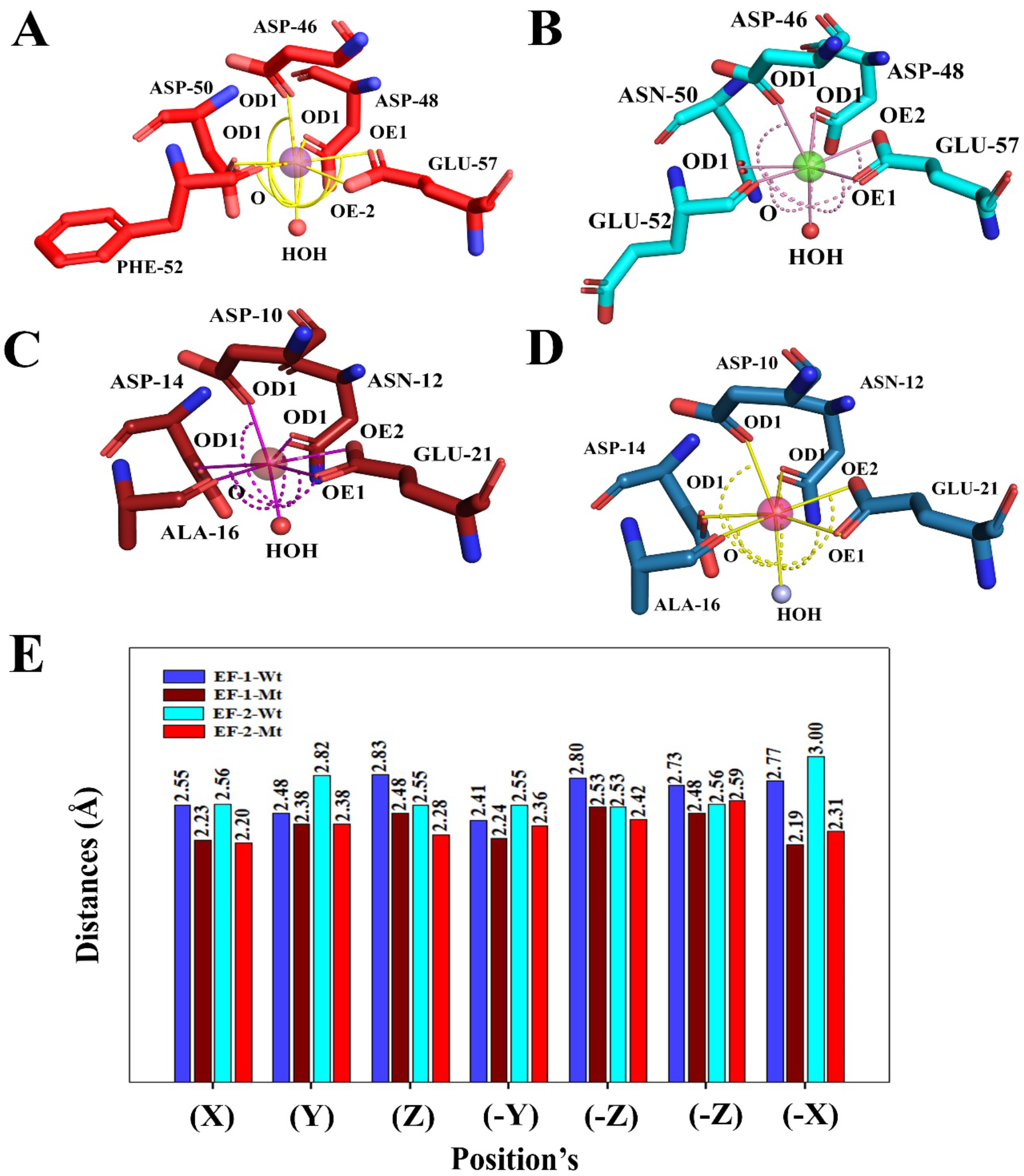
The arrangements of residues in the EF-hand Ca^2+^-bound sites of Nt*Eh*CaBP1-mutant and WtNt*Eh*CaBP1. A) The Ca^2+^-binding EF-2 site of the designed EhCaBP1-EF-2 mutant structure, showing the close contacts of the known ligands at the critical Ca^2+^-binding positions (X, Y, Z, -Y, -Z, -X) with the Ca^2+^ ion. The coordinating water molecule required for binding Ca^2+^ with a proper binding affinity was used to calculate various angles inside the cavity and to estimate the compactness of the site. B) The Ca^2+^-binding active site in the EF-2 loop of the Wt*Eh*CaBP1 protein. C) The EF1 loop of the Nt*Eh*CaBP1-EF2 mutant structure. D) The Ca^2+^-binding active site in the EF1 loop of the WtNt*Eh*CaBP1 structure. E) The distributions of distances from the oxygen atoms of the Ca^2+^-interacting residues to the Ca^2+^ ion for each of the two EF-hand sites, i.e., EF-1 and EF-2, of each of the two proteins, i.e., the WtNt*Eh*CaBP1 (blue) and Nt*Eh*CaBP1-mutant (red) proteins. For each of the four cases, these distances were calculated by averaging the distances of all the chains of the crystal structure. EF-1-Wt stands for the 1^st^ EF hand of the wildtype Nt*Eh*CaBP1, etc.

### The EF-1 loop of the Nt*Eh*CaBP1-EF2 mutant also shows reduced bond distance and bond angle

Although, we could not change any residues of EF-1 of Nt*Eh*CaBP1-EF2 mutant protein, but a comparative analysis of the Ca^2+^ coordinating distances of EF-1 of Nt*Eh*CaBP1-EF2 mutant structure also shows shrinkage in the Ca^2+^ coordinating sphere with respect to EF-1 of Wt-NtEhCaBP1 structure (Table 4) (Figure 6C and 6D). The OD1 atom of Asp-10, the first coordinating residue, was found to be positioned at the same site in both structures, but closer to the Ca^2+^ ion in the Nt*Eh*CaBP1-EF2 mutant than Wt-NtEhCaBP1 structure. Along with other residues which are involved in the Ca^2+^ coordination sphere, the 12^th^ position Glu-21 ligand was also observed to be closer to the Ca^2+^ ion in the mutant than in WtNt*Eh*CaBP1 (Table 4).

**Table 4.** List of the Ca^2+^-binding ligands at the conserved positions and water molecules of the Nt*Eh*CaBP1-mutant EF-1 loop and WtNt*Eh*CaBP1 EF-1-loop as shown in the figure 5 (C and D) along with the distances (Å) between the interacting atoms/ residues to the bound Ca^2+^ ion and the angles (Θ) with respect to the water molecule. Specifically listed are residues of EF-1 of chain A interacting with Ca^2+^ through their coordinating side-chain oxygens.

| Residue | AA | Angle ($\Theta$) | Position | Distance (d1) | Angle (Θ1) | Distance (d2) | Angle (Θ2) |
| --- | --- | --- | --- | --- | --- | --- | --- |
|  | **Binding Site (EF-I)** |  |  | ***EhCaBP1*-mutant EF-I** |  | ***EhCaBP1* EF I** |  |
| 10 | ASP | OD1-HOH-CA | **X** | **2.11** | 166.6 | 2.53 | 164.1 |
| 12 | ASN | OD1-HOH-CA | **Y** | **2.4** | 89.9 | 2.80 | 104.1 |
| 14 | ASP | OD1-HOH-CA | **Z** | **2.58** | 84.4 | 2.85 | 83.5 |
| 16 | ALA | O-HOH-CA | **-Y** | 2.25 | 96 | 2.43 | 89 |
| 21 | GLU | OE1-HOH-CA | **-Z** | 2.55 | 90.6 | 2.70 | 86.3 |
| 21 | GLU | OE2-HOH-CA |  | 2.46 | 93.5 | 2.70 | 107.6 |
| Water | HOH | NA | **-X** | 2.19 | NA | 2.80 | NA |

To overcome the problem of having small dataset, we included all six chains from two other reported structure of *Eh*CaBP1 (3LI6 and 2NXQ) and six chains of Nt*Eh*CaBP1-EF2 mutant structure (5XOP) for distance comparison of each Ca^2+^ ligating oxygen in EF-hand motifs (Figure 6E). The average Ca^2+^ coordination distance of NtEhCaBP1-EF2 mutant structure shows the shortest coordination distance for EF-2 than EF-1 and longest in EF1-of WtEhCaBP1 followed by EF-2 (Listings of the coordinating residues and distances for each of the individual chains from each structure are shown in Supplementary tables S7-S13).

### Cooperative binding enhances Ca^2+^ binding capability of the EF-1 loop of the mutant protein

EF-hand motifs mostly occur in pairs, and the Ca^2+^-binding mechanism and the resulting affinity also rely on how well these two motifs influence each other’s presence by interacting with the residues at the interface of Ca^2+^-binding sites. The non-covalent interactions play very important role in the stabilization of a pair of EF-hand loops [61, 64]. The communication between the two EF-hands in our structure was observed to be mediated by the non-covalent interactions involving conserved hydrophobic residues. We used the Residue Interaction Network Generator (RING) to identify covalent and non-covalent bonds, including π–π stacking and π–cation interactions, between the EF-1 and EF-2 motifs in both the Nt*Eh*CaBP1 and Nt*Eh*CaBP1-EF2 mutant structures (Figure 7) The Nt*Eh*CaBP1 structure showed numerous such interactions, including both at the C-terminal helix where F-58 and Y-62 formed π–π stacking interactions and at the binding interface where I-53 formed a hydrogen bond with V-17 (Figure 7A). In addition to a hydrogen bond between I-53 and V-17 as seen in the wildtype protein, an additional van der Waals interaction was observed between F-42 and I-45 (Figure 7B). The non-bonded interactions in the mutant were further enhanced by the incorporation of two additional phenylalanine (E52F and Q55F) in site 2. The analysis suggested that the more extensive hydrophobic interactions in the mutant helped its EF1 loop to move closer by 0.2 Å towards EF2 loop (supplementary figure S6). The detailed structural analysis using protein network nodes and edges as interactions suggested that the incorporation of the ring structures (of the aromatic residue) allowed the loops to interact through pi-pi interactions mediated via F-52 and F-55 and several aromatic-amide π interactions (Supplementary figure S6 D&E) and also increased the polarizability due to the face-to-face and edge-to-face interactions of aromatic rings exhibiting strong van der Waals interactions in the Nt*Eh*CaBP1 mutant.

**Figure 7.**
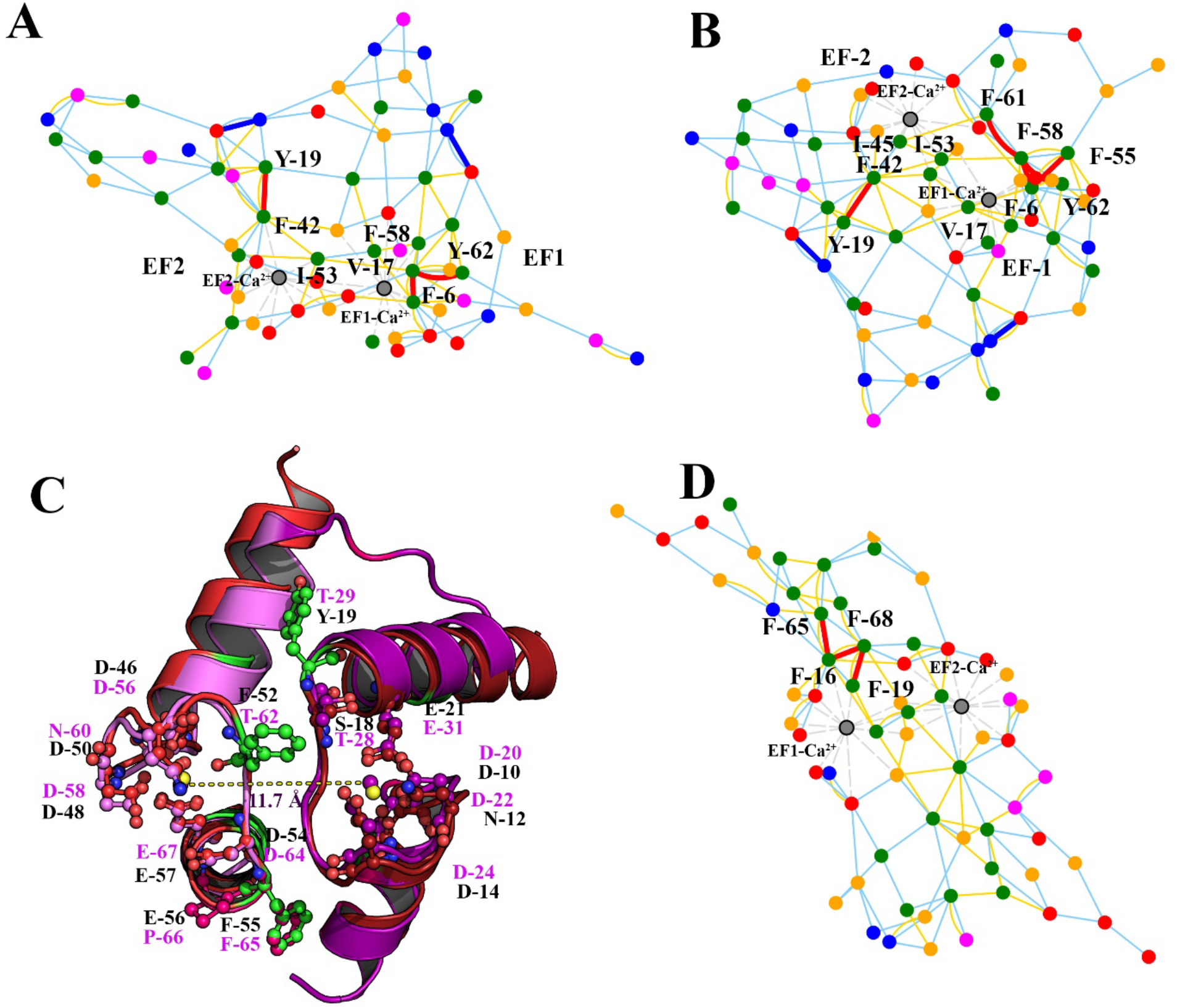
Structural basis of cooperativity of Ca^2+^ binding of WtNt*Eh*CaBP1(LAS), the Nt*Eh*CaBP1 mutant and CaM (HAS). Contact maps of A) WtNt*Eh*CaBP1 and B) the Nt*Eh*CaBP1 mutant where the residues at the interface between EF-1 and EF-2 are colored according to their physico-chemical properties: the hydrophobic amino acid residues are shown in green, positively charged residues in blue, negatively charged residues in red, polar residues in pink, and small residues such as alanine and glycine in yellow. The interactions are colored according to the type of bonding: hydrogen bonds are shown in cyan, van der Waals interactions in yellow, ionic interactions in blue, π–π stacking in red, and the metal-ion-mediated interactions in grey. The hydrophobic amino acid residues and the residues participating in π–π stacking are labelled. C) The two Ca^2+^-binding sites in the CaM (pink) and Nt*Eh*CaBP1 mutant (red) proteins, showing the positions of aromatic residues (shown in green color: Nt*Eh*CaBP1 mutant) mediating hydrophobic and van der Waals interactions. The residues forming the Ca^2+^-binding loop are labeled in both structures and the distance between the two-Ca^2+^ binding sites in CaM is also shown. D) The ionic interactions (blue), hydrogen bonds (cyan), and van der Waals interactions of the residues at the interface between EF-1 and EF-2.

### Insights from the high Ca^2+^ affinity of the calmodulin Ca^2+^-binding loop

In order to gain further insight into the different cooperative binding mechanisms exhibited by different pairs of EF-hand loops, we analyzed a high-Ca^2+^-affinity rat CaM (3SJQ) structure as a third model for comparison., since this CaM was found to be the closest CaM structural homolog (supplementary table S5) of the Nt*Eh*CaBP1 mutant. The CaM EF-1 (DKDGDGTITTKE) and EF-2 (DADGNGTIDFPE) shows SVM_Mar_ values of 0.93 and 1.87, respectively, and PSM_LogL_ scores of 5.58 and 6.39, respectively, hence both the sites were predicted a high affinity Ca^2+^ binder. The hydrophobic core formed by the interactions of two EF-hand motifs of CaM and Nt*Eh*CaBP1 EF-2 mutant were superimposed to understand what makes a strong Ca^2+^ binder. Surprisingly, the alignment showed a low RMSD of 0.1-0.2 Å. In CaM structure, the two Ca^2+^ ions of the two EF-hand motifs were 11.7Å away from each other (Figure 7C), compared to 11.8 Å for the Nt*Eh*CaBP1 mutant and 12Å for WtNt*Eh*CaBP1. Comparison of the contact map of CaM (Figure 7D) with NtEhCaBP1 EF-2 mutant shows the presence of more hydrophobic residues near the EF-1 motif of the high-affinity site (HAS) than low-affinity site (LAS). Analysis of the protein-protein interaction site in the Nt*Eh*CaBP1 EF-2 mutant shows eight aromatic residues compared to seven in CaM, i.e., with this difference due to the presence of Phe-52 in the Nt*Eh*CaBP1 EF-2 mutant but Thr-52 in CaM (Figure 7C). The close proximities of the two Ca^2+^-binding loops at the EF1-EF2 interface (Figure 7C) of CaM and Nt*Eh*CaBP1 EF-2 mutant suggested a stronger communication. The formation of an EFβ-scaffold in CaM and in the Nt-*Eh*CaBP1 EF-2 mutant apparently allows the sites to interact with each other through changes in the hydrogen bond network and increases the stability which is a major factor for high intrinsic binding affinity for Ca^2+^ [61].

### Dynamically coupled networks and correlated motions in EF-hand proteins

One of the factors that has been not yet been well studied in Ca^2+^-binding proteins is that how the motion of one domain of the protein influences that of the other and how the residues of these domains are networked together by long-range interactions so that they can manipulate each other’s functions by undergoing small conformational changes. During the comparative analysis throughout this study, we noticed dissimilarities of many side chain interactions, with these dissimilarities due to different orientations of the side chains (Supplementary Figure S4). The three model systems (Nt*Eh*CaBP1, the Nt*Eh*CaBP1 EF-2 mutant, and CaM) comprising six Ca^2+^-binding EF-hand motifs with different Ca^2+^-binding affinities were further analyzed to understand the role of the all the residues in influencing each other’s correlative movement. We calculated the dynamic cross-correlation (DCC) map and the dynamic protein residue network to understand the correlations amongst the residues in both a low-Ca^2+^-affinity protein (Nt*Eh*CaBP1) and high-Ca^2+^-affinity protein (Nt*Eh*CaBP1 mutant). Here, the protein structures were modeled as network systems by translating amino acid residues into nodes and both long- and short-range interactions into links (Figure 8). Comparing the full network of residues of Nt*Eh*CaBP1 (a monomer including EF1 and EF2), Nt*Eh*CaBP1 EF-2 mutant (a monomer including EF1 and EF2) and high Ca^2+^ affinity rat CaM (a monomer including EF2 and EF3). Interestingly this comparison revealed a pattern of communication leading to different correlated motions for the three proteins (Figures 8A, B and C). The community clustering map generated using the Girvan-Newman method showed the EF-hand motifs to be highly intramolecularly connected proteins with residues clustered as communities and numbered as clustered in the large networks. The Nt*Eh*CaBP1 with 65 residues, Nt*Eh*CaBP1 mutant with 66 residues, and N-terminal domain of CaM with 69 residues form the same set of nodes. Nt*Eh*CaBP1 forms 1527 edges, considerably more than the 1387 edges of the Nt*Eh*CaBP1 mutant and the 1397 edges of rat CaM. Both of the structures (i.e., Nt*Eh*CaBP1 mutant and CaM) with high-Ca^2+^-affinity binding sites showed a more compact structure as the clustering of intramolecularly connected communities resulted in fewer community nodes and edges than in WtNt*Eh*CaBP1 (supplementary table 11). The full network of Nt*Eh*CaBP1 formed the largest community3 (grey), comprising 23 residues (Figure 8A), whereas the high-Ca^2+^-affinity Nt*Eh*CaBP1 mutant formed a massive community 2 comprising 48 residues, quite similar to the CaM residue network that showed the largest community 2 with 38 residues. The differences in the communication between the clustered communities and the differences in the number of edges (connections) in the two very similar systems (Nt*Eh*CaBP1-mutant and Nt*Eh*CaBP1), i.e., differing by only 5 residues, was quite surprising. Six residues (Figure 8A) from the EF loops of Nt*Eh*CaBP1 were found to participate in the largest community (grey color) cluster compared to 13 residues in the Nt*Eh*CaBP1-mutant and 11 residues in CaM (community 2). The largest communities in each of the EF-hand motifs were determined to consist of residues mainly from the central helix. But the pattern varied in the high-affinity motifs, where the residues in the largest community were found to be distributed throughout the structure especially in the designed mutant (Figure 8B), all of whose C-terminal residues were observed to be part of the largest connected community. The high-Ca^2+^-affinity sites of CaM and Nt*Eh*CaBP1-mutant showed a high number of residues clustered together (Supplementary table S11). The determination of the number of nodes (residues) participating from individual domains shed light on the extent of the correlations amongst these 3 multiple EF-hand containing protein constructs. The N-terminal helix of NtEhCaBP1 showed residues connected to the largest community (grey), with the central helix having 13 residues and the C-terminal helix interconnected via 7 residues. The analysis of the high-affinity CaM and Nt*Eh*CaBP1-mutant proteins showed their N-terminal helices to have 13 and 12 residues connected to the largest community (red), their central helices to have 15 and 20 residues, and their C-terminal helices to have 9 and 15 residues, respectively. The comparison of these intra connected residues clearly suggested the extent of the direct correlation between the EF-loop site residues is higher in CaM than Nt*Eh*CaBP1, and highest in Nt*Eh*CaBP1-mutant.

**Figure 8.**
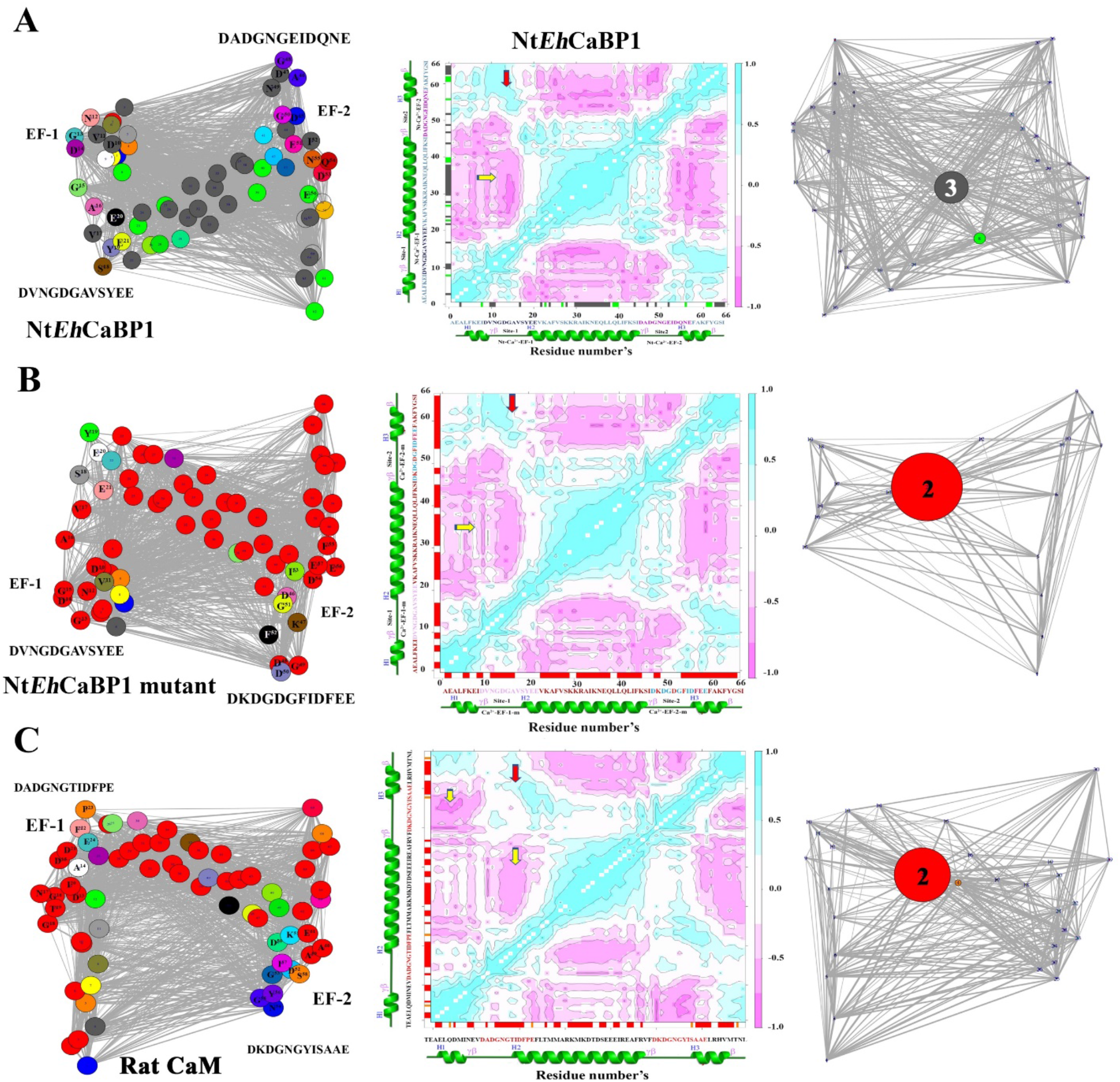
Comparison of the full networks of residues of three Ca^2+^-binding EF-hand proteins. The left column of images shows all-residue networks of monomers of A) Nt*Eh*CaBP1 B) Nt*Eh*CaBP1 mutant and C) N-terminal domain of CaM, with the amino acid residues (nodes) as circles and the long- and short-range interactions (links) as lines. Each network is colored by highly correlated local substructures (communities) within which the network connections are dense. The residues in the EF-loop are highlighted to show the position of the loop. The network communication in each 2D network comprises different communities colored and is shown using the force directed layout. The middle column of images shows the DCCM maps calculated by using the NMA ensemble with the Hessian matrix; the Cij value ranging from +1 to -1 are indicated by two colors, with cyan indicating positive correlation and pink indicating negative correlation amongst the residues. The leftmost column of images shows the community connections in each network. The rightmost column showing the largest communities numbered as per clustering are labeled in the image. The coarse grain network based on dynamically coupled communities shows the central helix residues forming the largest community connected with few residues from the N- and C-termini of Nt*Eh*CaBP1, in contrast to the Nt*Eh*CaBP1-mutant and CaM where the central helix is strongly connected to the residues from the EF loop site and the residues at the C-terminal end.

The dynamic correlation motions of the residues in EF-hand proteins plotted as DCC maps (figure 8) showed strong positive and negative correlation. Both WtNt*Eh*CaBP1 and Nt*Eh*CaBP1-mutant showed positive correlation in the EF1 loop, in contrast to that in the EF2 loop (Figure 8). The pattern of correlation shown by the EF-hand proteins suggested a direct correlation between the two Ca^2+^-binding sites as shown by the red arrow in the DCC map (Figure 8). The yellow arrow shows the negative correlation between the central helix and the EF loops (Figure 8). In summary, the dynamic correlation and the clustering analysis suggested that the coordination between two EF-hand motifs varies in high & low binding, in high binders it resulted in strong coordination of residues. All of the EF loops directly influence the central helix by long-and short-range interactions affecting the residue network thereby enhancing the allosteric interactions.

## Discussion

In this study we upgraded our earlier designed software ‘Calbinder’ in a way so that along with identification of an EF-hand protein sequence user can also design and manipulate Ca^2+^ binding affinity of their protein of interest, as well. To confirm that our upgraded software helps in manipulating the Ca^2+^ binding affinity accurately or reaching to our expectation, we designed a high affinity EF-hand motif sequence and validated our hypothesis by exploiting biophysical and structural approach. As it was predicted, the newly designed EF-hand motifs of NtEhCaBP1-EF2 mutant protein yielded a higher Ca^2+^ binding affinity (ITC data) than the wild type protein. Further, with respect to Wt-NtEhCaBP1, the EF-2 of NtEhCaBP1-EF2 structure, shows a more compact Ca^2+^ coordination sphere, suggest the tight binding of Ca^2+^. The compact Ca^2+^ coordination sphere induced structural bend in the helix-3, due to which helix-3 of one trimer interact with the helix-2 of another trimer leading to the hexamer formation in the solution as well as in the crystal state. We anticipate that the incorporated mutation in NtEhCaBP1-EF-2 protein induced the cooperativity, i.e., binding of Ca^2+^ at one site induced the binding at second site. Due to the induced cooperative in Ca^2+^ binding, the NtEhCaBP1-EF-2 form of protein may binds Ca^2+^ with higher affinity.

As, the crystal structure of Wt-NtEhCaBP1 has Ca^2+^ bound at its both EF-hand motifs [28], therefore the observation of week Ca^2+^ binding in ITC experiment of Nt-EhCaBP1 was a surprising observation. A justified statement for this concern is that the crystallization trials of Wt-NtEhCaBP1 was performed in the presence of high Ca^2+^ concentration (5mM), which allows Ca^2+^ ion to accommodate in the crystal lattice, therefore it observed in the crystal structure. Further, to check if higher concentration of Ca^2+^ may allow better binding for Wt-NtEhCaBP1 and the binding curve may achieve the saturation in ITC experiment, we stepwise increased the amount of Ca^2+^ till 25mM, however it did not help in achieving the saturation point similar or close to Nt-EhCaBP1 EF-2 mutant form of protein (data not shown). Suggesting that Wt-Nt-EhCaBP1 form of protein could not bind Ca^2+^ with high affinity even in presence high Ca^2+^ concentration.

Binding affinity of Ca^2+^ to EF-hand motifs containing CaBPs is widely governed by the residues present in EF-hand loop and the Ca^2+^ binding affinity can be manipulated usually by altering the 1^st^, 3^rd^, 5^th^, and 12^th^ residues of the active site. Altering these critical Ca^2+^ binding residues of EF-hand motifs may drastically reduce the Ca^2+^ binding affinity or completely abolish the binding of Ca^2+^. Such approach reduces the possibility study the function of these proteins over a wide range of Ca2+ binding affinity. Our software addresses this issue and one can design/manipulate the EF-hand motif sequence to achieve the desired Ca^2+^ binding affinity, as per their experimental requirement. Most importantly with this software, manipulation of Ca^2+^ binding affinity is not limited only with the residues which are directly involved in Ca^2+^ coordination rather one can explore all the twelve residues of the EF-hand motif and manipulate the Ca^2+^ binding affinity by performing permutation combination of entire twelve residues present at EF-hand loop. Such flexibility may allow user to manipulate the Ca^2+^ binding affinity without affecting the conserve structural core (EF-hand motifs) of CaBPs. Although the experimental data presented in this manuscript is from only one protein, nevertheless it provides the strong confidence of machine learning based method of Ca^2+^ binding affinity manipulation.

With enough data, we expect it to be possible to factor the participation of cooperative binding into a predictive model that should provide deeper insight into the physiological Ca^2+^-binding state. In summary, the non-bonded interactions at the interface of two EF loops and the ionic interactions at the Ca2+-binding sites changed the binding characteristics of EF1 in the designed protein despite the residues of EF1 not having been altered. The allosteric communication from the coupling of structural changes at the Ca2+-binding sites helped regulate the binding characteristic of the proteins. As an allosteric effector, the EF2 mutation was shown to modulate the behavior of the EF1 site. This form of intramolecular communication provided insight into the role of residues that participates in Ca2+ binding via dynamically coupled coordination with other residues in the structure. We expect the structure-function relationship obtained from the analysis of EF hand proteins to aid the building of highly accurate predictive models of Ca^2+^ binding in EF hand proteins. We also expect a better understanding of the various interactions carried out by all of the residues, i.e., not only those at the active sites but the entire EF hand motifs, to help in building machine-learned models capable of predicting the effect of cooperativity. Moreover, the understanding of accurate protein networks generated from high-resolution structural data can further enable the designing of similar proteins mimicking the EF-hand motif functions.

